# Anterior expansion and posterior addition to the notochord mechanically coordinate embryo axis elongation

**DOI:** 10.1101/2021.01.20.427467

**Authors:** Susannah B.P. McLaren, Benjamin J. Steventon

## Abstract

During development the embryo body progressively elongates from head-to-tail along the anterior-posterior (AP) axis. Multiple tissues contribute to this elongation through a combination of convergence and extension and/or volumetric growth. How force generated by the morphogenesis of one tissue impacts the morphogenesis of other axial tissues to achieve an elongated axis is not well understood. The notochord, a rod-shaped tissue possessed by all vertebrates, runs across the entire length of the somitic compartment and is flanked on either side by the developing somites in the segmented region of the axis and presomitic mesoderm in the posterior. Cells in the notochord undergo an expansion that is constrained by a stiff sheath of extracellular matrix, that increases the internal pressure in the notochord allowing it to straighten and elongate. Therefore, it is appropriately positioned to play a role in mechanically elongating the somitic compartment. Here, we use multi-photon mediated cell ablation to remove specific regions of the developing notochord and quantify the impact on axis elongation. We show that anterior notochord cell expansion generates a force that displaces notochord cells posteriorly relative to adjacent axial tissues and contributes to the elongation of segmented tissue during post-tailbud stages of development. Crucially, unexpanded cells derived from progenitors at the posterior end of the notochord provide resistance to anterior notochord cell expansion, allowing for force generation across the AP axis. Therefore, notochord cell expansion beginning in the anterior, and addition of cells to the posterior notochord, act as temporally coordinated morphogenetic events that shape the zebrafish embryo AP axis.

## Introduction

The formation of complex body shapes is a central question in developmental biology. A key component of this process is the elongation of the embryo head-to-tail, or anterior-posterior (AP) axis. Axis elongation is brought about by the physical deformation of multiple axial tissues as they undergo morphogenesis (Bénazéraf et al., 2017; Steventon et al., 2016, Xiong et al., 2020). The vertebrate embryo body axis is segmented into blocks of tissue called somites, that later form the skeletal muscle and vertebrae of the adult body (Gomez et al., 2008). Running through the middle of the embryo, the rod-shaped notochord is flanked on either side by the somitic compartment: the somites in the segmented region of the axis and the presomitic mesoderm in the posterior (Stemple, 2005; Oates et al., 2012).

Disrupting the morphogenesis of the notochord results in axis truncation and severe defects such as skeletal malformations (Bagwell et al., 2020; Ellis et al., 2013). Studies in Xenopus embryos have identified the circumferentially constrained expansion of cells within the notochord to lead to an increase in notochord stiffness as the embryo develops, making it a candidate for driving the physical deformation of surrounding somitic tissue that may be required for AP axis elongation (Adams et al., 1990). Notochord cells also expand in zebrafish embryos as they develop fluid-filled vacuoles (Ellis et al., 2013), and measurements of volumetric tissue growth have revealed an increase in notochord volume as development progresses (Steventon et al., 2016). However, it remains unclear how force generated by notochord morphogenesis may impact the elongation of the somitic compartment and extend the zebrafish embryo AP axis.

Concomitant with notochord cell expansion, a separate morphogenetic event occurs in which cells are contributed to the posterior end of the notochord from a population of notochord progenitors (Kanki and Ho, 1997). Early termination of progenitor addition to the notochord is associated with AP axis elongation defects (Row et al., 2016). Whilst the role of other tailbud-located progenitor populations in elongating the AP axis has been investigated (Goto et al., 2017; Mongera et al., 2018; Attardi et al., 2018), it is not known how the posterior contribution of cells from notochord progenitors is coordinated with the expansion of more anterior notochord cells, and how these combined processes impact axis elongation.

As notochord morphogenesis progresses somites are segmented from the unsegmented presomitic mesoderm in the posterior tailbud until 30 to 34 somites are formed in total, with the tailbud remaining present until the 30-somite stage when it begins to degenerate (Oates et al., 2012; Kimmel et al., 1995). As somites mature, they undergo bending until a ‘chevron shape’ is reached (Rost et al., 2014; Tlili et al., 2019). Whilst elongation generated from unsegmented tailbud-derived tissue has been studied (Das et al., 2019; Dray et al., 2013; Lawton et al., 2013; Mongera et al., 2018; Steventon et al., 2016), how elongation progresses in mature segmented tissue during post-tailbud stages of development is less clear.

Here, we utilise targeted multi-photon tissue ablation to investigate the physical impact of notochord morphogenesis on the surrounding somitic compartment, and identify the concomitant expansion of notochord cells, and contribution of progenitors to the posterior end of the notochord, as two temporally coordinated morphogenetic events that shape the zebrafish embryo AP axis.

## Results and Discussion

### Modes of axis elongation change as embryos progress from tailbud stages into post-tailbud stages of development

To quantitatively characterise embryo AP axis elongation occurring concomitantly with notochord volume increase, we live-imaged developing zebrafish embryos from the onset of notochord cell expansion and quantified elongation in segmented and unsegmented regions of the axis (Fig. 1A and movie 1). Throughout this study we use the somitic compartment as a reference for investigating axis elongation and utilize somite boundaries to demarcate different regions of the axis. Segmented tissue elongation is tracked using the boundaries of 5 formed somites. Unsegmented tissue length is defined using the posterior-most somite boundary and tail-end, and the length generated from this region incorporates newly formed somites as development progresses.

**Fig. 1.**
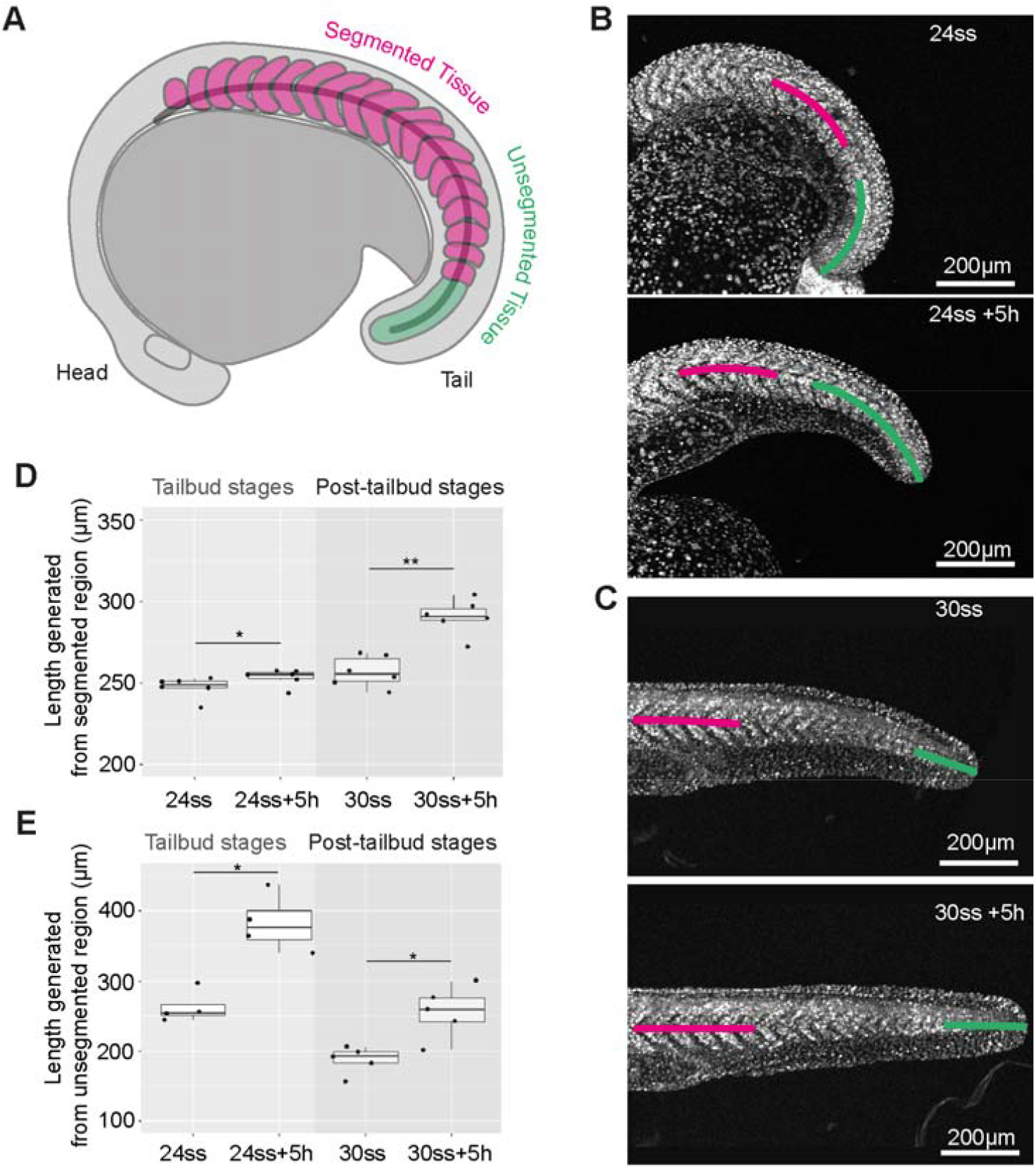
Axis elongation occurs predominantly in the unsegmented region during tailbud stages and later continues in segmented tissue. (A) Schematic outlining the segmented and unsegmented regions of the somitic compartment. (B) A live zebrafish embryo expressing NLS-KikGR at the 24-somite stage and 5 hours later. (C) A live zebrafish embryo expressing NLS-KikGR at approximately the 30-somite stage and 5 hours later. (D) The length of a region of segmented tissue in tailbud (24ss) and post-tailbud (30ss) embryos and length generated from this region over 5-hours (n=6 and n=6 respectively, p<0.05, p<0.01 respectively). (E) The length of a region of unsegmented tissue in tailbud (24ss) and post-tailbud (30ss) embryos and length generated from this region over 5-hours (n=4 and n=5 respectively, p<0.05, p<0.05 respectively). Curves for calculating segmented (magenta) and unsegmented tissue (green) length are overlaid. Anterior is to the left and posterior to the right in all images.

We compared elongation over a 5-hour period during tailbud (24ss) and post-tailbud stages (30ss) to investigate the modes of axis elongation utilised during each of these phases of embryo morphogenesis. Segmented tissue elongation was minimal during tailbud stages (Fig.1B and D magenta region), with a mean length of approximately 6µm (2.5% increase) generated in the measured region (Fig.1D). However, in post-tailbud stage embryos mean length increased by approximately 34µm (13% increase) (Fig.1C and E). In contrast, the mean length generated from unsegmented tissue (Fig.1B green region) was approximately 120µm (45% increase) in tailbud stage embryos, and 70µm (36% increase) in post-tailbud stage embryos (Fig.1E). When measuring the length of the whole somitic compartment we found the increase in mean length to be lower than expected based on previous studies (Kimmel et al., 1995) (Fig.S1A and B). This is likely due to a difference in the way axis length is measured, with our measurements accounting for changes in embryo curvature.

To investigate embryo straightening we measured the average curvature of the somitic compartment as well as the trunk and tail regions of the axis (Fig. S1A). Embryos underwent a dramatic straightening, with curvature decreasing in all regions (Figs.S1C, D and E). Overall, our findings show that embryo AP axis elongation can be described by two phases: straightening of the embryo, accompanied by elongation generated from unsegmented posterior tissue during tailbud stages, followed by elongation of segmented tissue during post-tailbud stages (movie 1).

### Notochord cell expansion leads to the posterior displacement of notochord cells relative to adjacent axial tissues

To gain insight into how notochord morphogenesis may be impacting axis elongation during tailbud and post-tailbud stages of development, we characterised notochord cell expansion by measuring the AP length of vacuoles in live embryos both temporally, in an equivalent region of the notochord of different stage embryos (Fig.2A), and spatially, in notochord regions in the anterior vs. the posterior in embryos of the same stage (Fig.2C). In agreement with previous work, we observed that notochord cells expanded over time (Ellis et al., 2013; Bagwell et al., 2020) (Fig.2B), and that vacuoles in more anterior regions were longer than those in more posterior regions (Fig.2D). In embryos entering post-tailbud stages, we found that notochord cells began to move posteriorly relative to neighbouring axial tissues, in contrast to the bi-directional displacement of notochord cells recently observed in Medaka (Seleit et al., 2020) (Fig. 2E and F, movie 2). Manual tracking of more expanded notochord cells in the anterior and less expanded cells in the posterior revealed that the posterior displacement of notochord cells relative to adjacent tissues was greater in more expanded regions (Figs. S2A and B). These findings suggest that notochord cell expansion generates a force that leads to the posterior displacement of notochord cells relative to adjacent axial tissues.

**Fig. 2.**
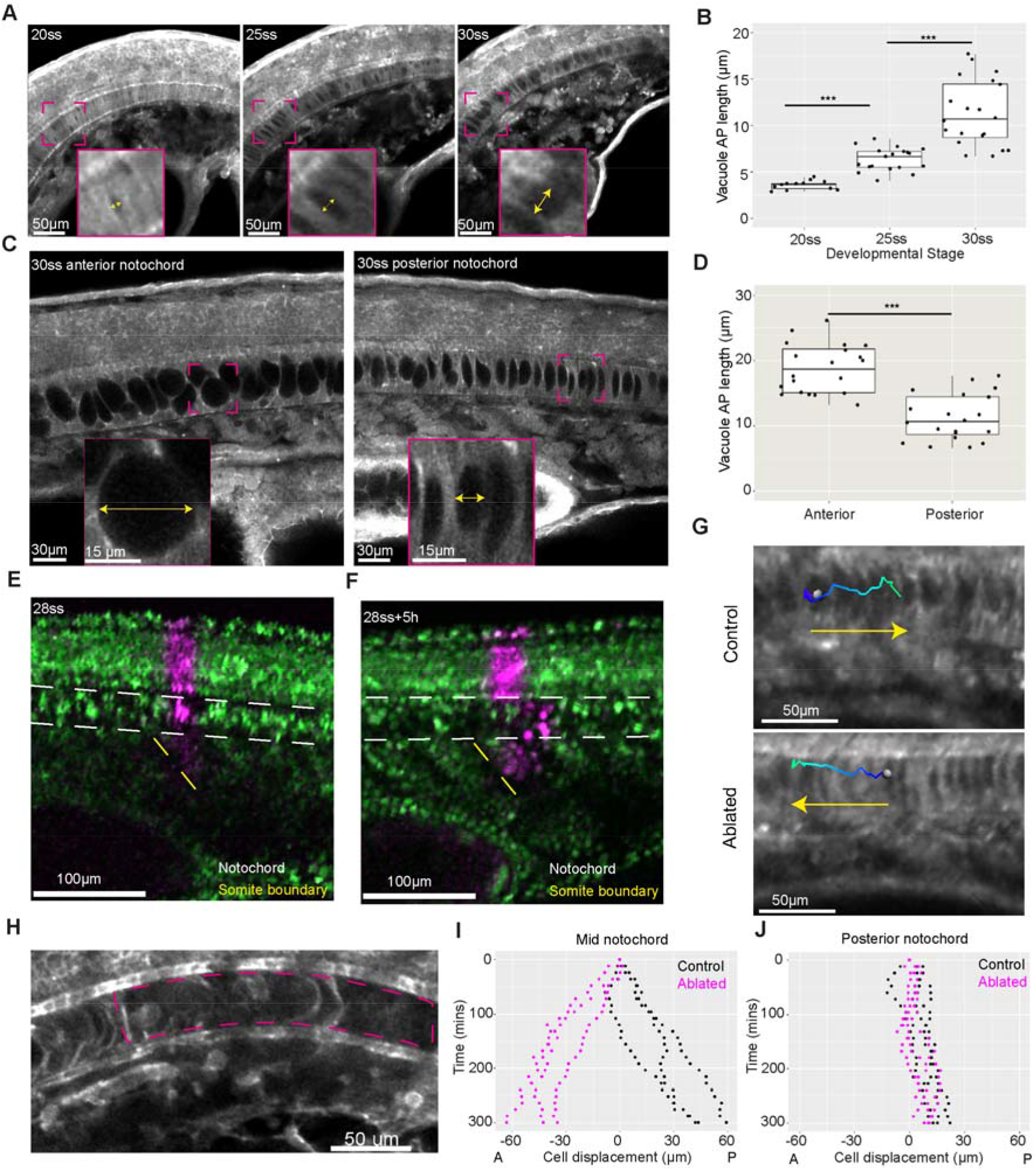
Notochord cell expansion leads to the posterior displacement of notochord cells relative to adjacent axial tissues. (A) Notochord cells in an equivalent region of a Lifeact-GFP expressing embryo at 20, 25, and 30-somite stages. Yellow arrows indicate direction in which length measurements were taken. (B) AP length of vacuoles in µm (n=3, n=4 and n=4 embryos per stage respectively, 3 to 5 vacuoles measured per embryo. p<0.001 and p<0.001 respectively). (C) Notochord cells in the anterior and posterior of a Lifeact_GFP expressing embryo. Yellow arrows indicate direction in which length measurement was taken. (D) AP length of vacuoles in µm in anterior and posterior regions of 30-somite stage embryos (n= 4 embryos, 5 vacuoles measured per region. p<0.001). (E) A midline view of a 28-somite stage embryo expressing NLS-KikGR showing a photolabel made in the neural tube, notochord, and somites at the same region of the axis. (F) A midline view of a 28-somite stage embryo expressing NLS-KikGR showing the photolabel in (E) 5 hours later. (G) Expanding notochord cell tracks in control and ablated embryos. (H) A region of ablated notochord imaged approximately 3 hours after ablation in a Lifeact-GFP expressing embryo. (I) Cell displacement in µm of manually tracked expanded notochord cells in control and ablated embryos (n=3 embryos per condition, 1 track per embryo). (J) Cell displacement in µm of manually tracked unexpanded notochord cells in control and ablated embryos (n=3 embryos per condition, 1 track per embryo). Anterior is to the left and posterior to the right in all images.

In order to investigate how forces generated from anterior notochord cell expansion might propagate along the AP axis and to neighbouring tissues, a technique is required that allows for precise spatio-temporal removal of notochord cells. To this end, we used multi-photon ablation to cut through notochord cells at defined regions of the AP axis (movie 3), resulting in the destruction of tissue local to the ablation site (Fig. 2H). Ablation of anterior notochord cells reversed the direction of displacement of expanding notochord cells posterior to the ablation site (Fig.S2C and movie 4), whilst the movement of unexpanded posterior notochord cells was less affected (Figs. 2G, I and J). Thus, notochord cell expansion progresses posteriorly along the notochord in a correlated fashion with straightening and tail elongation. As embryos enter post-tailbud stages the continued expansion of cells in the notochord generates a force, displacing neighbouring expanded notochord cells relative to adjacent somites.

### Notochord cell expansion is a driver of segmented tissue elongation during post-tailbud stages of development

Elongation during tailbud stages was driven by the segmentation of posterior unsegmented tissue. To investigate whether notochord cell expansion impacts on this mode of elongation, we performed site-specific ablations of anterior notochord cells just as they were beginning to expand and measured posterior tail elongation during tailbud stages of development (Fig.3A and S3A). Posterior tail elongation was not significantly affected in ablated embryos (Fig.3B and C). Strikingly, segmentation-derived elongation could continue in an embryo with almost complete notochord ablation (movie 5), and presomitic mesoderm progenitors continued to exit the tailbud in ablated embryos (Fig.3D). To investigate the impact of notochord cell expansion on the straightening of the axis, we measured the curvature of a 15-somite sample of the axis in control and ablated embryos over time (Fig.S3B). We found that embryo straightening was perturbed in regions where the notochord was ablated (Fig.S3C) suggesting that notochord continuity is required for complete straightening of the axis, whilst straightening appeared to continue in regions where the notochord was intact. Together, these results demonstrate that elongation derived from the segmentation of posterior unsegmented tissue is a robust process that can continue in the absence of notochord cell expansion.

**Fig. 3.**
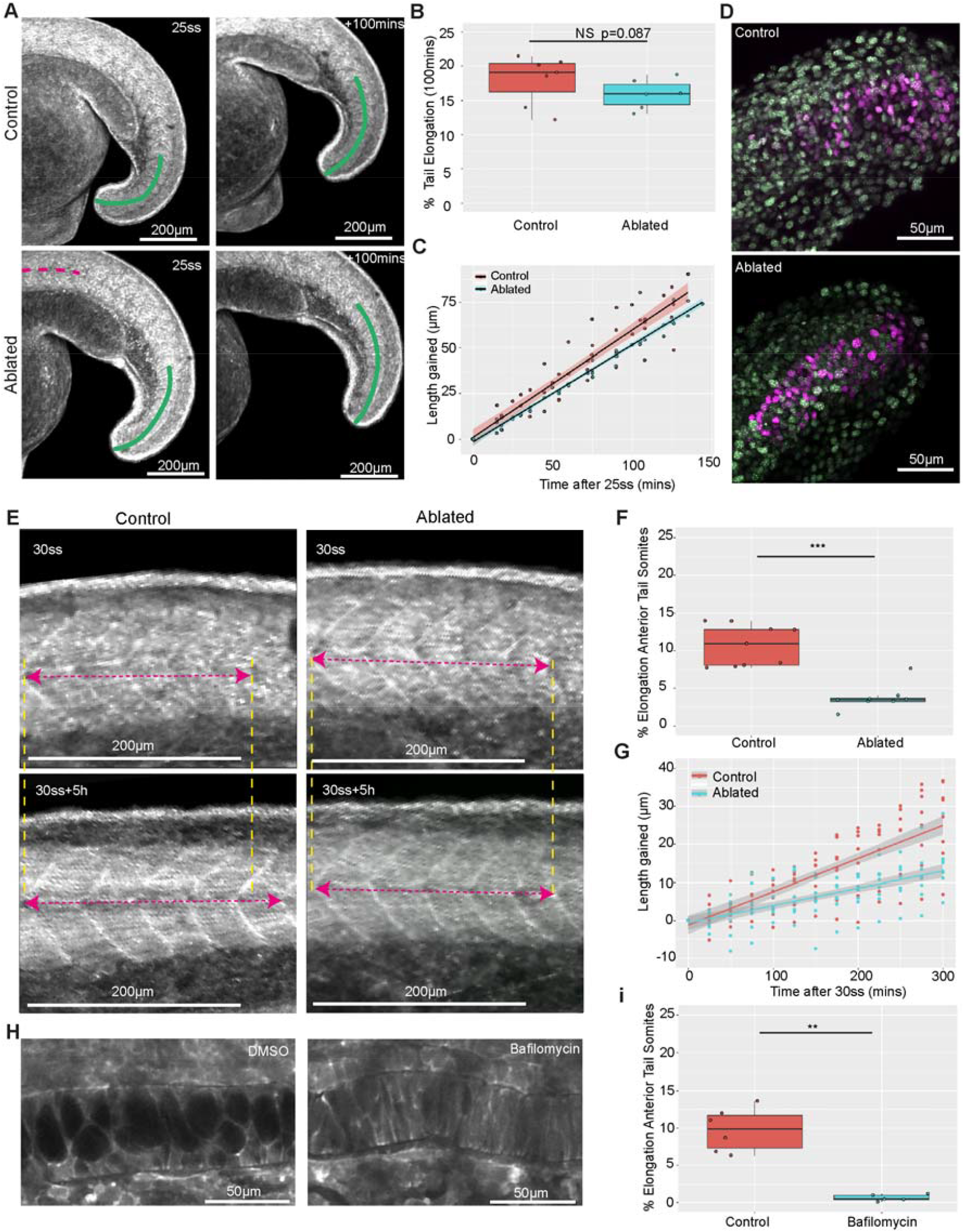
Notochord cell expansion contributes to segmented tissue elongation during post-tailbud stages. (A) Control and ablated Lifeact-GFP expressing embryos imaged at the 25-somite stage and 100 minutes later. Overlaid curves show the region tracked for elongation measurements. The magenta line indicates the region where the notochord is ablated. (B) Percentage elongation of the tail region in control and ablated embryos (n=7 control embryos, n=6 ablated embryos, p=0.087). (C) Length gained in the posterior tail of tailbud stage control and ablated embryos over time. (D) Photolabels of presomitic mesoderm progenitors in control and ablated embryos imaged 4 hours after the time of labelling. (n=4 control and n=5 ablated embryos). (E) Elongation of a 5-somite region in control and ablated post-tailbud stage Lifeact-GFP expressing embryos. (F) Percentage elongation of a 5-somite region in control and ablated post-tailbud stage embryos over a period of 5 hours (n=9 control and n=8 ablated embryos, p<0.001) (G) Length gained in a 5-somite region in control and ablated post-tailbud stage embryos over time. (H) Images of a notochord DMSO and Bafilomycin treated Lifeact-GFP expressing embryos. (I) Percentage elongation of a 5-somite region in control and Bafilomycin treated embryos over 3 hours (n=6 control and n=5 Bafilomycin treated embryos, p<0.01). Anterior is to the left and posterior to the right in all images.

To investigate whether notochord cell expansion impacts the segmented tissue elongation observed during post-tailbud stages, we measured the length of a 5-somite region from the 30-somite stage over a 5-hour period in embryos with and without anterior notochord ablations (Fig. 3E). The region used for quantifying segmented tissue elongation was located in an equivalent region along the axis in controls and ablated embryos and positioned posteriorly away from the site of notochord ablation in the ablated case. As characterised above, this region was adjacent to either posteriorly directed notochord cell movement (in controls) or anteriorly directed notochord cell movement (in ablated embryos). Segmented tissue elongation was approximately halved in ablated embryos (Figs.3F and G, movies 6 and 7), and inhibiting notochord cell expansion by blocking vacuolation with Bafilomycin (an inhibitor of vacuolar type H^+^-ATPases required for vacuole expansion) (Ellis et al., 2013) also reduced the elongation of segmented tissue (Figs.3H, I, and S3D). Together these results reveal a role for notochord cell expansion in elongating segmented tissue in post-tailbud stage embryos.

### Notochord progenitors provide a source of resistance to anterior notochord cell expansion that facilitates the elongation of segmented tissue

We hypothesised that notochord cell expansion elongated segmented tissue via an AP oriented stretch. This would require notochord cell expansion to be resisted in the AP direction. We observed that unexpanded notochord cells in the posterior exhibited less movement relative to adjacent somites than more anterior expanded cells, suggesting that they may resist the force generated by notochord cell expansion. We wondered if the addition of unexpanded cells to the posterior notochord was coordinated with the progression of notochord cell expansion posteriorly along the axis. To investigate this, we used the expression of *flh* to mark the notochord progenitor domain and measured the volume of the domain at different stages of development (Fig. S4A). We found that notochord progenitor domain volume decreased as development progressed (Fig. S4C) and that this corresponded to a decrease in the length contributed to the posterior end of the notochord by progenitors (Fig. S4B and D). Next, we attempted to prevent the contribution of cells to the posterior notochord by ablating the notochord progenitor population (Fig. S4E). Whilst axis elongation defects occurred (Fig. S4F), the notochord progenitor pool was able to regenerate after ablation and recover the posterior notochord (Figs. S4H and I). Intriguingly, elongation resulting from the segmentation of the final few somites in the region where the posterior notochord had been recovered was unaffected (Figs. S4G and J). These findings highlight the robustness of posterior processes that contribute to axis elongation, both in terms of cell contribution to the posterior notochord and segmentation-derived elongation.

Given that the notochord regenerated after notochord progenitor ablation, we decided to focus our perturbations on the already formed posterior notochord. In embryos entering post-tailbud stages of development notochord cell expansion has progressed posteriorly along the axis to the anterior tail region (close to the end of the yolk extension) where it meets the unexpanded notochord cells (Fig.4D). To investigate whether unexpanded notochord cells are subject to a force generated by anterior notochord cell expansion, we imaged the nuclei of unexpanded notochord cells in this region in control embryos, and just posterior to an ablation site in ablated embryos (Fig. 4A). Nuclei in control embryos were angled towards the posterior, whereas nuclei posterior to an ablation site were angled towards the anterior (Fig.4B and C), suggesting that unexpanded notochord cells are being deformed by a posteriorly directed force arising from notochord cell expansion.

**Fig. 4.**
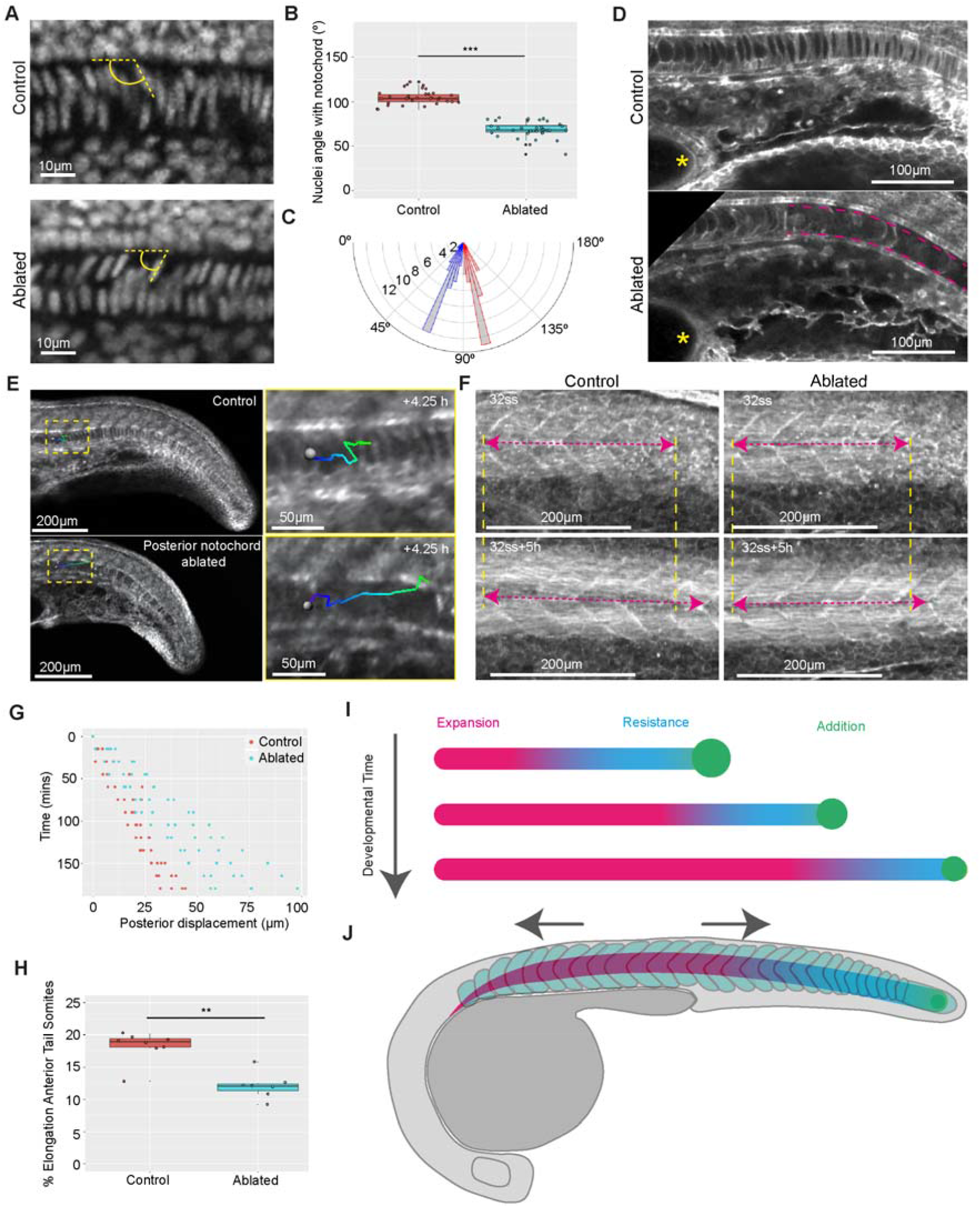
Notochord progenitors provide a source of resistance to notochord cell expansion, facilitating segmented tissue elongation. (A) DAPI-stained notochord nuclei located in the anterior tail region in fixed control and ablated embryos. (B) Angles between dorsally located nuclei and the notochord AP axis (º) in control and ablated embryos (n=7 embryos per condition, 5 angles measured per embryo, p<0.001). (C) A polar histogram showing the distribution of angles between dorsally located nuclei and the notochord AP axis (º) in control and ablated embryos (n=7 embryos per condition, 5 angles measured per embryo, p<0.001). (D) Notochord cells located close to the end of the yolk extension (yellow asterisk) in a Lifeact-GFP expressing control embryo and an embryo with a posterior notochord ablation (magenta lines). (E) Manual tracking of expanding notochord cells in control embryos and embryos with a posterior notochord ablation. (F) Elongation of a 5-somite region in control and posteriorly-ablated post-tailbud stage embryos. (G) Notochord cell displacement over time in control and posteriorly-ablated embryos (n=4 and n=5 embryos respectively). (H) Percentage elongation of a 5-somite region in control and posteriorly-ablated embryos (n=8 and n=7 embryos respectively, p<0.01). (I) Notochord cell expansion progresses posteriorly along the axis and is resisted by unexpanded notochord cells contributed by posterior notochord progenitors. (J) Notochord cell expansion generates an AP stretch along the somitic compartment. Anterior is to the left and posterior to the right in all images.

To investigate whether unexpanded posterior notochord cells provided a source of resistance to notochord cell expansion, we ablated these cells in embryos at post-tailbud stages (Fig. 4D) and manually tracked expanding notochord cells just anterior to the ablation site (Fig. 4E). The posterior displacement of expanding cells was increased in ablated embryos compared to controls (Fig. 4G, movies 8 and 9). In addition, the elongation of a region of somites anterior to the ablation site (in an equivalent region to that used in previous ablation experiments) was decreased in ablated embryos (Figs. 4F and H). These findings suggest that posterior unexpanded notochord cells provide a source of resistance to notochord cell expansion that may facilitate stretching of the AP axis, deforming segmented tissue in post-tailbud stage embryos.

Our findings show that notochord cell expansion generates a force which is resisted by unexpanded notochord cells that are derived from posterior notochord progenitors (Fig. 4I). Given the posterior progression of notochord cell expansion along the axis, and the depletion of notochord progenitors, we expect that the region resisting notochord cell expansion will shrink as development progresses, until the entirety of the notochord is in an expanded state. Indeed, experiments in which the tail tip of larval-stage embryos has been cut away reveal a rapid extrusion of notochordal tissue, suggesting that the posterior notochord no longer provides resistance at these stages (Romero et al., 2018). Somite elongation is decreased in embryos in which notochord cell expansion and force propagation along the notochord has been disrupted, leading us to propose that this force is translated into an AP oriented stretch, elongating segmented tissue in post-tailbud stage embryos (Fig. 4J). This may be facilitated by physical coupling between the notochord and somitic compartment in the posterior of the embryo and uncoupling in the trunk (Dray et al., 2013; Tlili et al., 2019).

Investigation into multi-tissue mechanical interactions in avian embryos has shown that, in contrast to the zebrafish, the presomitic mesoderm expands in volume leading to the lateral-to-medial compression of the notochord, displacing the tip of the notochord posteriorly into the tailbud and facilitating progenitor contribution to the axis (Bénazéraf et al., 2017; Steventon et al., 2016; Xiong et al., 2020). Our findings suggest that the notochord in zebrafish embryos contributes to axis elongation via a different mechanism that depends on the progressive expansion of notochord cells. Studies approximating the notochord as a hydraulic skeleton have shown that such a system is capable of exerting force on its surroundings (Koehl et al., 2000). We find that notochord cell expansion generates a force that deforms segmented tissue during post-tailbud stages of development, contributing to AP axis elongation.

Whether this mechanism of axis elongation is utilised in other embryo species during later stages of development is an area for future investigation.

## Acknowledgements

We thank Andrea Dimitracopoulos, Fengzhu Xiong and Bertrand Benazeraf for critical reading of the manuscript, and members of the Steventon, Martinez-Arias and Franze group meetings for their support and feedback on the project. We also thank the PDN fish facility for their unwavering help and support with animal care and maintenance, and the Cambridge Advanced Imaging Centre for their support with laser ablation experiments. B.S. was supported by a Henry Dale Fellowship jointly funded by the Wellcome Trust and the Royal Society (109408/Z/15/Z) and S.B.P.M. was supported by the Wellcome Trust funded Developmental Mechanisms PhD programme.

## Author Contributions

S.B.P.M. and B.S. conceived the project. S.B.P.M. carried out experiments and analysed the data. S.B.P.M. and B.S. wrote the manuscript.

## Materials and Methods

### Zebrafish strains and maintenance

This research was regulated under the Animals (Scientific Procedures) Act 1986 Amendment Regulations 2012 following ethical review by the University of Cambridge Animal Welfare and Ethical Review Body (AWERB).

Adult Zebrafish (Danio rerio) and embryos were reared at 28°C and staged using the number of somites. Wildtype embryos used in this study were of the TL strain. Prior to live imaging embryos were anaesthetised with tricaine (MS-222).

The following zebrafish lines were used in this study; H2B-GFP, Tg(actb2:Lifeact-EGFP), and pNtl:Kaede^16^.

### Timelapse Live Imaging

Embryos were mounted in 3% Methylcellulose to hold embryos still whilst allowing movement of the tail and ensuring embryos could continue to develop normally.

Embryos were mounted in glass bottom dishes (MatTek), with the head and yolk submerged in methylcellulose but with the tail freed of methylcellulose. Embryos were submerged in E3 media and an eyelash tool was used to reposition embryos in the methylcellulose so that they were lying flat on the bottom of the dish. Dishes with mounted embryos were transferred to a Zeiss LSM 700 confocal microscope equipped with a heated chamber set to 28°C. A 10x air objective (NA=0.45) was used to capture the whole body of developing embryos for timelapse movies.

### Drug treatments

Bafilomycin A1 (Sigma Aldrich) was used to inhibit vacuolation in the notochord of developing zebrafish embryos. 16ss embryos were submerged in E3 media with Bafilomycin at a final concentration of 0.5uM. Embryos were then incubated for 6 hours at 28°C. Vacuolation in the notochord of Lifeact-GFP embryos was imaged live using a 20x air objective. Live imaging of somite elongation in DMSO and Bafilomycin treated embryos was performed using pNtl:kaede-expressing embryos pre-treated with 0.25µM Bafilomycin (or DMSO of an equivalent volume) and imaged from the 30-somite stage using a multi-well chamber.

### Photolabelling

Wild-type Zebrafish embryos were injected with approximately 200 pg of nuclear-targeted Kikume at the one-cell stage and incubated in the dark. Embryos were mounted and transferred to the microscope as described above for live imaging. For notochord cell movement analysis, rectangular stripes were defined along the embryo. Photoconversion of NLS-KikGR was performed whilst using a 20x air objective and scanning the 405 nm laser at 11% power within defined regions. Photolabelling was confirmed by simultaneously visualing both NLS-KikGR emissions using the 488 nm and 561nm lasers.

### Cell tracking

The centre of notochord cells in lifeact-GFP expressing embryos was manually tracked in imaris relative to a nearby somite boundary located in an equivalent region of the axis in each embryo. Reference frames were generated using the ‘reference frame’ tool, the x axis was aligned with the AP axis and the origin was placed at the somite boundary.

### Hybridisation Chain Reaction (HCR)

HCR was used to visualise gene expression in fixed embryos. HCR has previously been described here (Choi et al., 2014). The notochord progenitor domain was visualised using probes targeted to the gene *flh* (also known as *noto*).

### Photoablation

Laser ablation was performed using a TriM Scope II Upright 2-photon scanning fluorescence microscope equipped with a tunable near-infrared laser. Inspector Pro software was used to control the microscope and a laser power of approximately 1.3W was used for ablation.

Notochord ablations were carried out using a lifeact-GFP reporter line to visualise notochord cells and ablations were performed in the plane of the notochord.

For notochord progenitor ablations the morphology of the notochord progenitor cell population was used to determine the location of the notochord progenitors.

Ablations were carried out to ensure as many of the notochord progenitors as possible were ablated, without ablating any other progenitor types.

### Image processing and analysis

Images were processed either in ImageJ/Fiji or Imaris.

### Length and curvature measurements

For length and curvature measurements the ImageJ plug-in ‘Kappa’ was used. NLS-KikGR, H2B-GFP and Lifeact-GFP expressing embryos were used for length and curvature measurements. A maximum projection of the nuclear-GFP or actin-GFP channel was used so that somite boundaries could be visualised and used to define the segmented and unsegmented regions. Embryo side views were used to measure the curvature and length of regions of the axis. Trunk curvature was measured using points to generate a spline, and the points were adjusted so that a smooth curve fitting the curvature of the region passed through the midpoint of somites in that region. Tail curvature was measured in a similar fashion. The average curvature was used for comparing the curvature of these regions in multiple embryos of different stages. To visualise the curvature profile along the whole somitic compartment in embryos of different stages, a spline passing from the anterior tip to the posterior tip of the somitic compartment was generated, and point curvature was used to generate a heatmap in order to visualise the curvature profile along the axis.

### Notochord progenitor domain volume

To measure the volume of the notochord progenitor domain at different developmental stages *flh* (also known as *noto)* gene expression was visualised using HCR and acquisition of confocal z-stacks of fixed 20ss, 25ss, and 30ss embryo tails. These images were imported into imaris, allowing for visualisation in 3D. The ‘surfaces’ function in imaris was used to generate a surface around the *flh* gene expression signal, with surface volume given as an output.

### Vacuole AP length measurements

The ‘oblique slicer’ tool in imaris was used to find a plane that passed through the middle of the notochord where the maximum length of vacuoles could be properly visualised. The ‘measurements’ tool was used to make two points per vacuole defining a line passing through the greatest AP length of the vacuole. Line lengths were calculated using an inbuilt function in imaris. Vacuole lengths were measured in wildtype or DMSO treated Lifeact-GFP expressing embryos.

### Nuclei angle measurements

The ‘measurements’ tool in imaris was used to measure the angle between nuclei located dorsally in the notochord and the dorsal AP edge of the notochord in fixed DAPI stained embryos with and without notochord ablations.

### Data analysis

Data was stored in excel spreadsheets and analysis was performed using the programming language ‘Python’. All data plots were generated in Python using the ggplot library. Box plots show the median, upper and lower quartiles, and whiskers represent 1.5 times the interquartile distance. All data points were shown and included in statistical analyses.

### Statistical Analysis

The Mann-Whitney U nonparametric statistical test was used to test whether two independent samples came from populations with the same distribution. This test was implemented in python using the scipy.stats.mannwhitneyu() function.

The Kruskal-Wallis nonparametric statistical test was used to test whether more than two independent samples came from populations with the same distribution.

Pairwise comparisons between samples were performed in a post-hoc fashion to identify whether compared samples came from populations with the same distribution. The Kruskal-Wallis test was implemented using the stats.kruskal() function, and the post hoc test was implemented with the sp.posthoc_conover() function.

In all cases a p-value of <0.05 was taken as a threshold for significance.

## Supplementary information

**Fig. S1.**
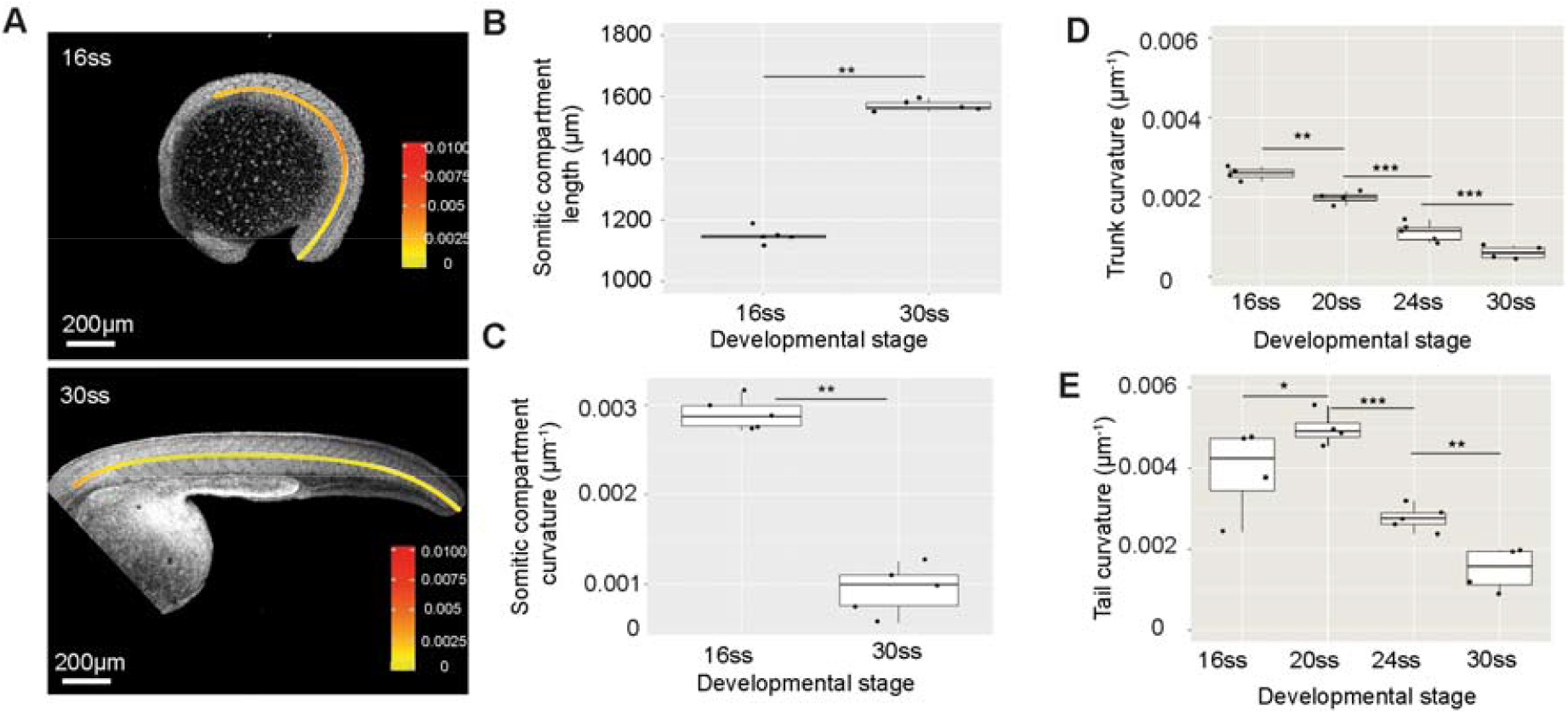
Elongation and straightening during tailbud stages of development. (A) Representative images of live 16 and 30 somite stage embryos. Overlaid curves indicate region used for somitic compartment length and curvature measurements. Heatmap shows point curvature. (B) Somitic compartment length in 16 and 30 somite stage embryos (n=5 per stage, p<0.01). (C) Somitic compartment length in 16 and 30 somite stage embryos (n=5 per stage, p<0.01). (D) Trunk curvature in 16, 20, 24 and 30 somite stage embryos (n=4,4,5,4 embryos respectively per stage, p<0.01, p<0.001, p<0.001 respectively). (E) Tail curvature in 16, 20, 24 and 30 somite stage embryos (n=4,4,5,4 embryos respectively per stage, p<0.05, p<0.001, p<0.01 respectively).

**Fig. S2.**
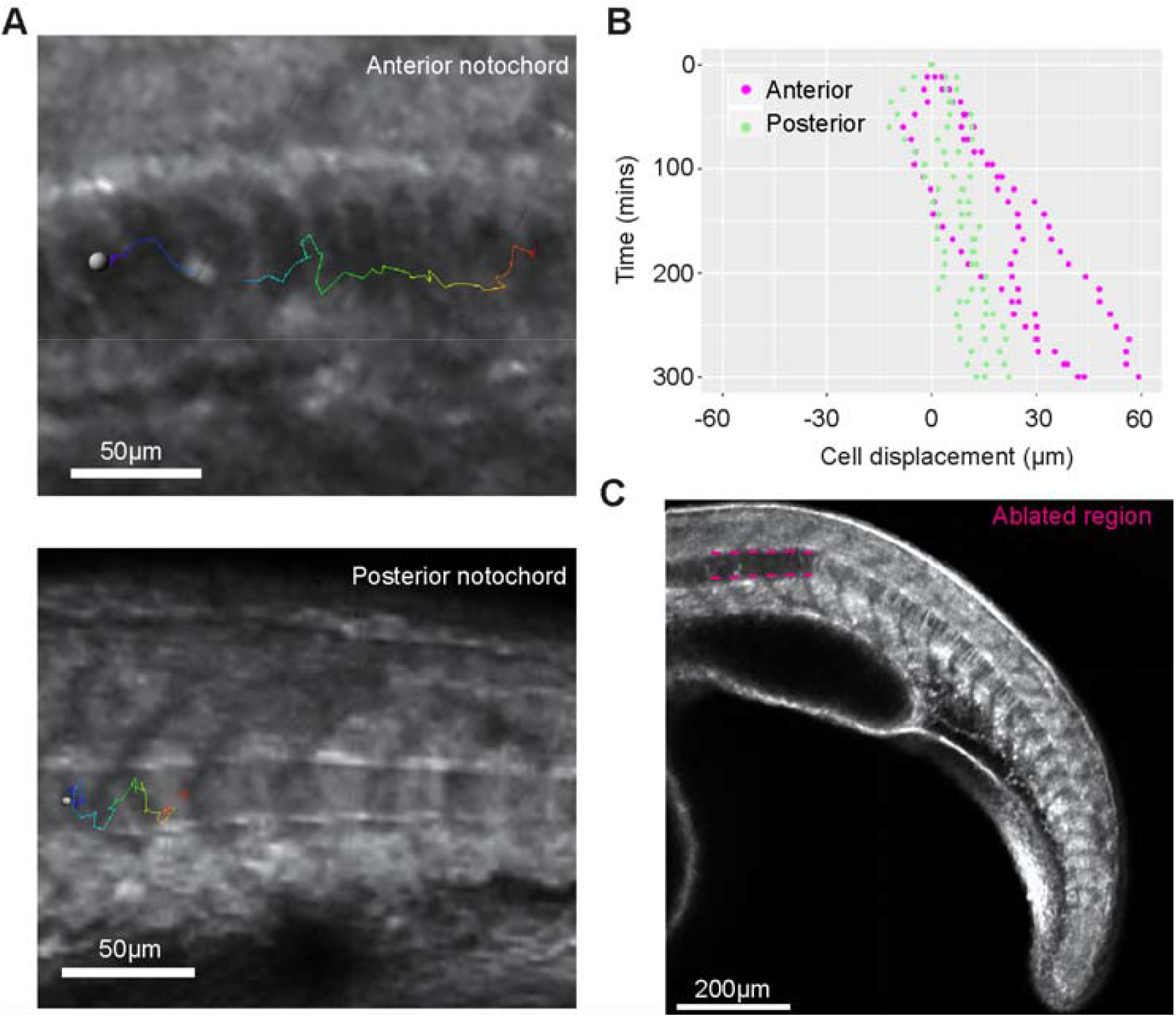
Notochord cell displacement relative to adjacent somites in expanded and unexpanded notochord regions. (A) Notochord cell tracks in an anterior expanded region and a posterior unexpanded region. (B) Notochord cell displacement over time in expanded trunk regions and unexpanded tail regions (n= 3 embryos). (C) An anterior notochord ablation in a lifeact-GFP expressing embryo.

**Fig. S3.**
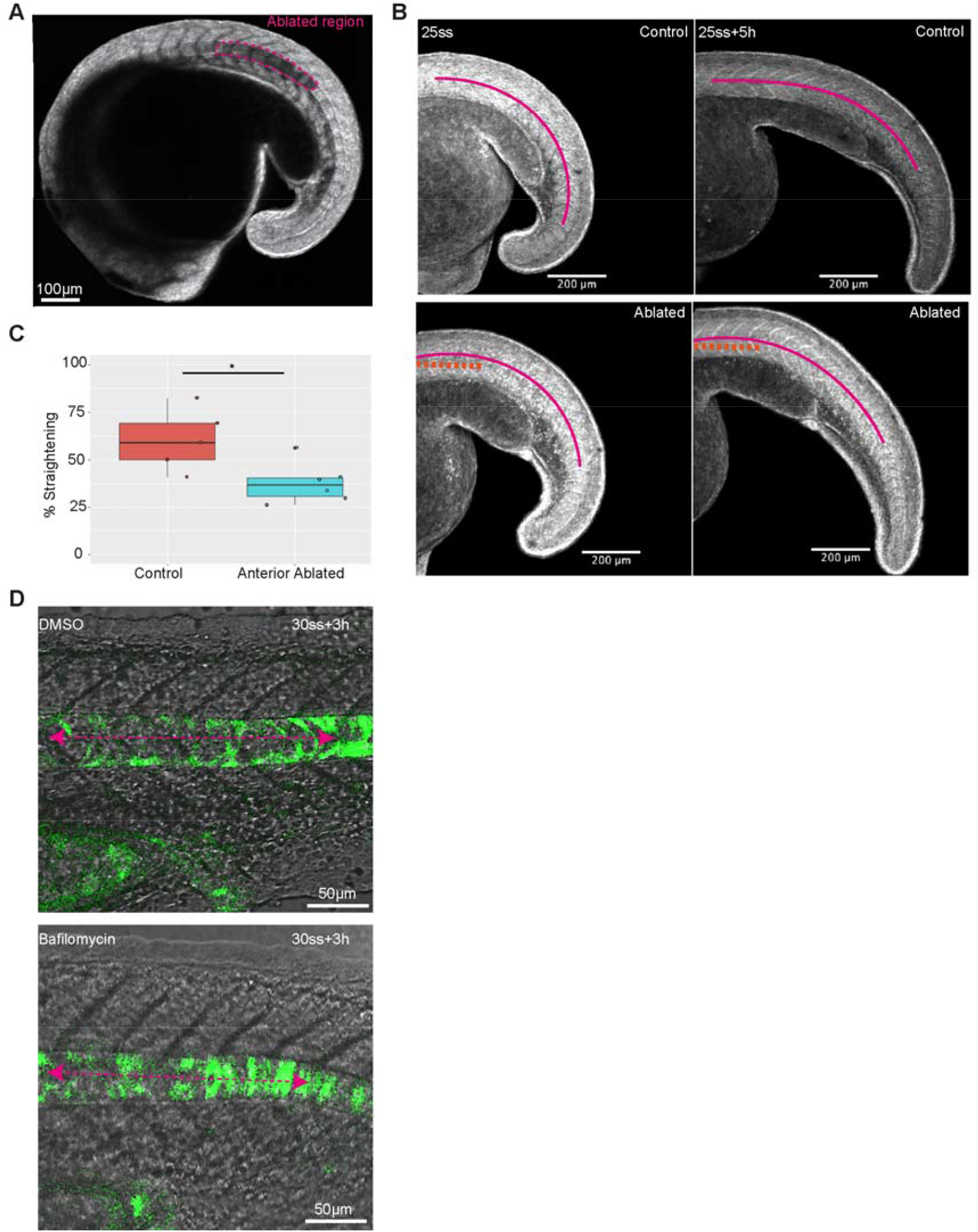
Notochord cell expansion impacts embryo straightening and elongation. (A) Representative image of an embryo with a region of anterior notochord ablated just as vacuoles are beginning to expand. (B) Images of control and anterior notochord ablated embryos at the 25 somite stage and 5 hours later. Overlaid curves show the region tracked over time for curvature measurements. (C) Percentage straightening (decrease in curvature) over 5 hours in control and ablated tailbud stage embryos (n=5 control and 6 ablated embryos, p<0.05). (D) A region of 5 anterior tail somites in control and bafilomycin treated embryos imaged 3 hours after the 30 somite stage.

**Fig. S4.**
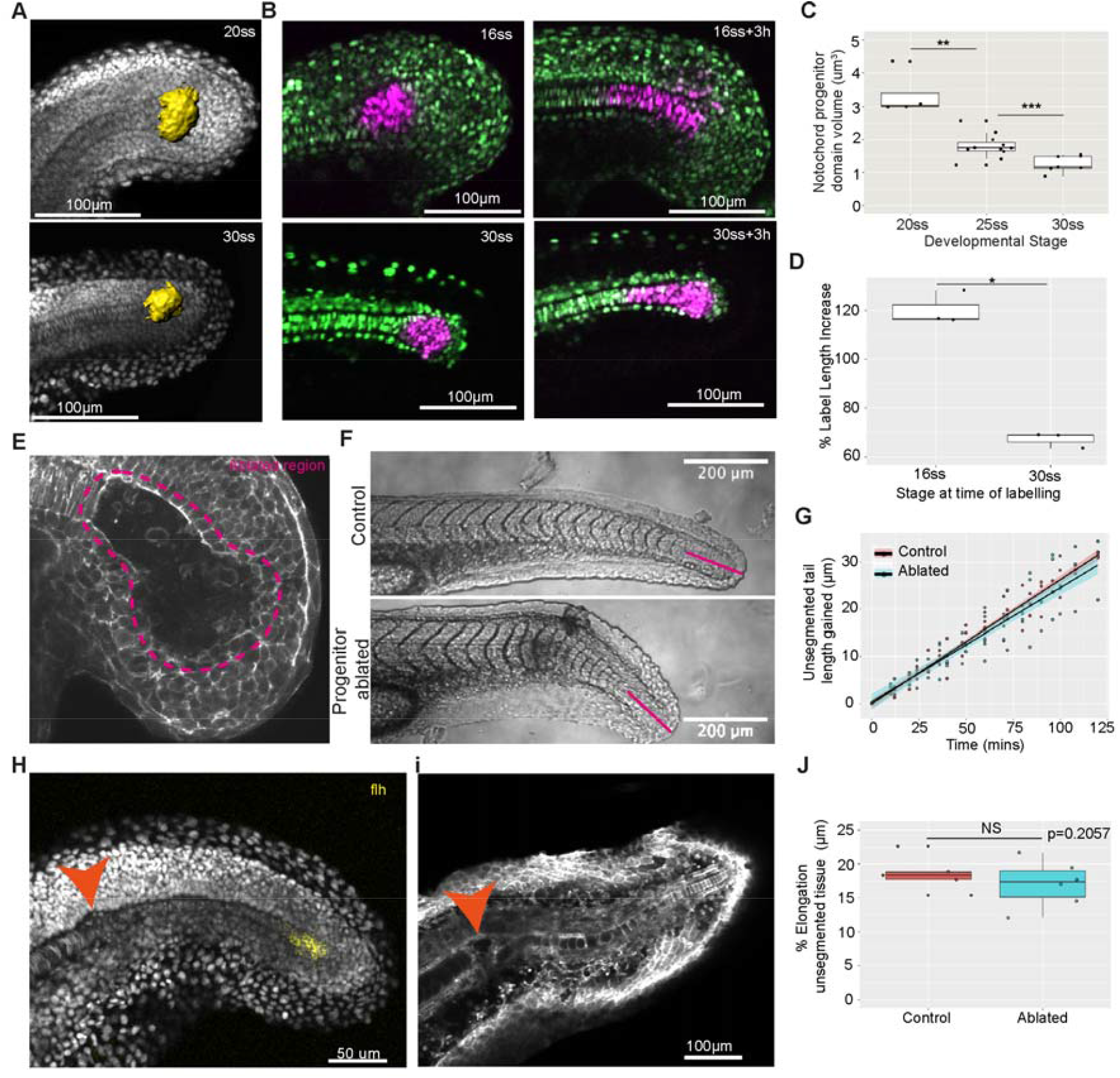
Notochord progenitors regenerate after ablation and recover the posterior notochord. (A) 3D surfaces of the notochord progenitor domain generated using *flh* gene expression in 20 and 30 somite stage embryos. (B) Notochord progenitor photolabels in 16 and 30 somite stage embryos with corresponding images taken 3 hours after labelling. (C) Notochord progenitor domain volume in µm^3^ in 20, 25 and 30 somite stage embryos (n=5, n=13, n=7 embryos respectively, p<0.01, p<0.001 respectively). (D) Photolabel AP length increase over 3 hours in 16 and 30 somite stage embryos (n=3 embryos per stage, p<0.05). (E) Representative image of a 16-somite stage embryo with notochord progenitors ablated. (F) Representative images of control and progenitor ablated post-tailbud stage embryos. The magenta line indicates the length of a small region of residual unsegmented tissue tracked over time. (G) Length gain in a small region of unsegmented tissue over time in control and progenitor ablated post-tailbud stage embryos (n=6 control and n=7 ablated embryos). (H) Representative image of an embryo fixed approximately 5 hours after notochord progenitor ablation. The gene expression marker ‘flh’ marks the notochord progenitors. (I) Representative image of an embryo imaged approximately 24 hours after notochord progenitor ablation. The recovered notochord is visualised using a lifeact-GFP reporter. (J) Percentage elongation of unsegmented tissue in the tail region after recovery from notochord progenitor ablation over approximately 100 minutes (n=5 control and n=6 ablated embryos, p=0.2057). Orange arrows indicate the internalised ablated tissue.

